# Orthogonal targeting of SAC1 to mitochondria implicates ORP2 as a major player in PM PI4P turnover

**DOI:** 10.1101/2023.08.28.555163

**Authors:** Colleen P. Doyle, Andrew Rectenwald, Liz Timple, Gerald R. V. Hammond

## Abstract

Oxysterol binding protein (OSBP)-related proteins (ORPs) 5 and 8 have been shown to deplete the lipid phosphatidylinositol 4-phosphate (PI4P) at sites of membrane contact between the endoplasmic reticulum (ER) and plasma membrane (PM). This is believed to be caused by transport of PI4P from the PM to the ER, where PI4P is degraded by an ER-localized SAC1 phosphatase. This is proposed to power the anti-port of phosphatidylserine (PS) lipids from ER to PM, up their concentration gradient. Alternatively, ORPs have been proposed to sequester PI4P, dependent on the concentration of their alternative lipid ligand. Here, we aimed to distinguish these possibilities in living cells by orthogonal targeting of PI4P transfer and degradation to PM-mitochondria contact sites. Surprisingly, we found that orthogonal targeting of SAC1 to mitochondria enhanced PM PI4P turnover independent of targeting to contact sites with the PM. This turnover could be slowed by knock-down of soluble ORP2, which also has a major impact on PM PI4P levels even without SAC1 over-expression. The data reveal a role for contact site-independent modulation of PM PI4P levels and lipid antiport.

## Introduction

The exchange of lipids by non-vesicular lipid transport is a core function of membrane contact sites between organelles (Prinz et al., 2020). This process can be mediated by soluble or tethered lipid transfer proteins that bind and solubilize individual lipids, or else proteinaceous conduits that permit lipids to flow between membranes (Wong et al., 2019). However, these proteins present a thermodynamic conundrum: Lipid exchange is known to enrich specific lipids in one membrane relative to the other at the contact site, yet the proteins themselves employ what is essentially a passive transport process. In other words, how can a lipid transfer protein move a lipid up a concentration gradient?

A seminal discovery shed light on one answer to this question (Saint-Jean et al., 2011): the discovery that Osh4p, a yeast member of the ORP family, can bind to both sterols and a phosphoinositide, PI4P. It followed that ORPs could employ an anti-porter mechanism; specifically, a phosphoinositide concentration gradient is created between the trans-Golgi network (TGN) and the ER by TGN localized PI 4-kinases (PI4Ks) that generate PI4P, and the ER-localized SAC1 phosphatase that degrades it. Transport of PI4P down this gradient by Osh4p releases energy used to counter-transport sterol from the ER back to the TGN, up its gradient. Ultimately, the energy of ATP hydrolysis is coupled to sterol transport via PI4P turnover. The mechanism was subsequently demonstrated for cholesterol transport at the mammalian TGN by OSBP (Mesmin et al., 2013), and was expanded to include PS transport from ER to PM by ORP5 and ORP8 in humans (Chung et al., 2015) and the homologues Osh6p and Osh7p in yeast (Moser von Filseck et al., 2015).

An implicit assumption of this model is that SAC1 activity acts to deplete PI4P and maintain the gradient, acting downstream of PI4P transport. It follows that ablation of SAC1 activity would lead to the accumulation of PI4P in the ER, which has been observed (Cheong et al., 2010; Zewe et al., 2018). However, PI4P transport by ORP family proteins remains controversial. An alternative hypothesis posits that ORP proteins act as negative regulators of PI4P signaling by sequestering the lipid, and that this inhibition can be relieved by high local concentrations of the counter lipid such as sterol (Wang et al., 2019b). Notably, such a model does not specifically require localization of the ORP at a membrane contact site with the ER. Instead, the alternative lipid cargoes modulate available PI4P pools at the same membrane. Differentiation between sequestration verses transport mechanisms can be difficult in cells, where both hypotheses predict decreases in accessible PI4P levels by ORPs in donor membranes like the PM or TGN.

In this paper, we attempt to answer the question: To what extent does PM PI4P catabolism depend on lipid transport to the ER via membrane contact sites? Our approach was to reconstitute the PI4P transfer and hydrolysis steps at orthogonal, PM:mitochondria contact sites. Surprisingly, we find that simply re-targeting SAC1 to the mitochondrial outer membrane, independent of membrane contact sites, is sufficient to accelerate PM PI4P turnover. The data implicate the non-contact site targeting ORP2 as a major mediator of PM PI4P turnover via non-vesicular lipid transport.

## Results

To test mechanisms of PM PI4P catabolism in cells, we began with experiments to engineer the most potent PM PI4P lipid transfer protein identified to date, ORP5 (Chung et al., 2015; Sohn et al., 2018). ORP5 contains an N-terminal extended Pleckstrin Homology (PH) domain that interacts selectively with the PM via PI4P and PI(4,5)P_2_ (Sohn et al., 2018); this is connected to the ORD lipid-transfer domain by a long region of low complexity, and is anchored to the ER by a C-terminal transmembrane helix (cartoon in **Fig. 1A**). When expressed as an mCherry-fusion, ORP5 localizes to ER-PM contact sites, colocalizing with the ER-PM contact site marker MAPPER (Chang et al., 2013) when viewed by equatorial or ventral confocal sections (**Fig. 1A**).

**Figure 1:**
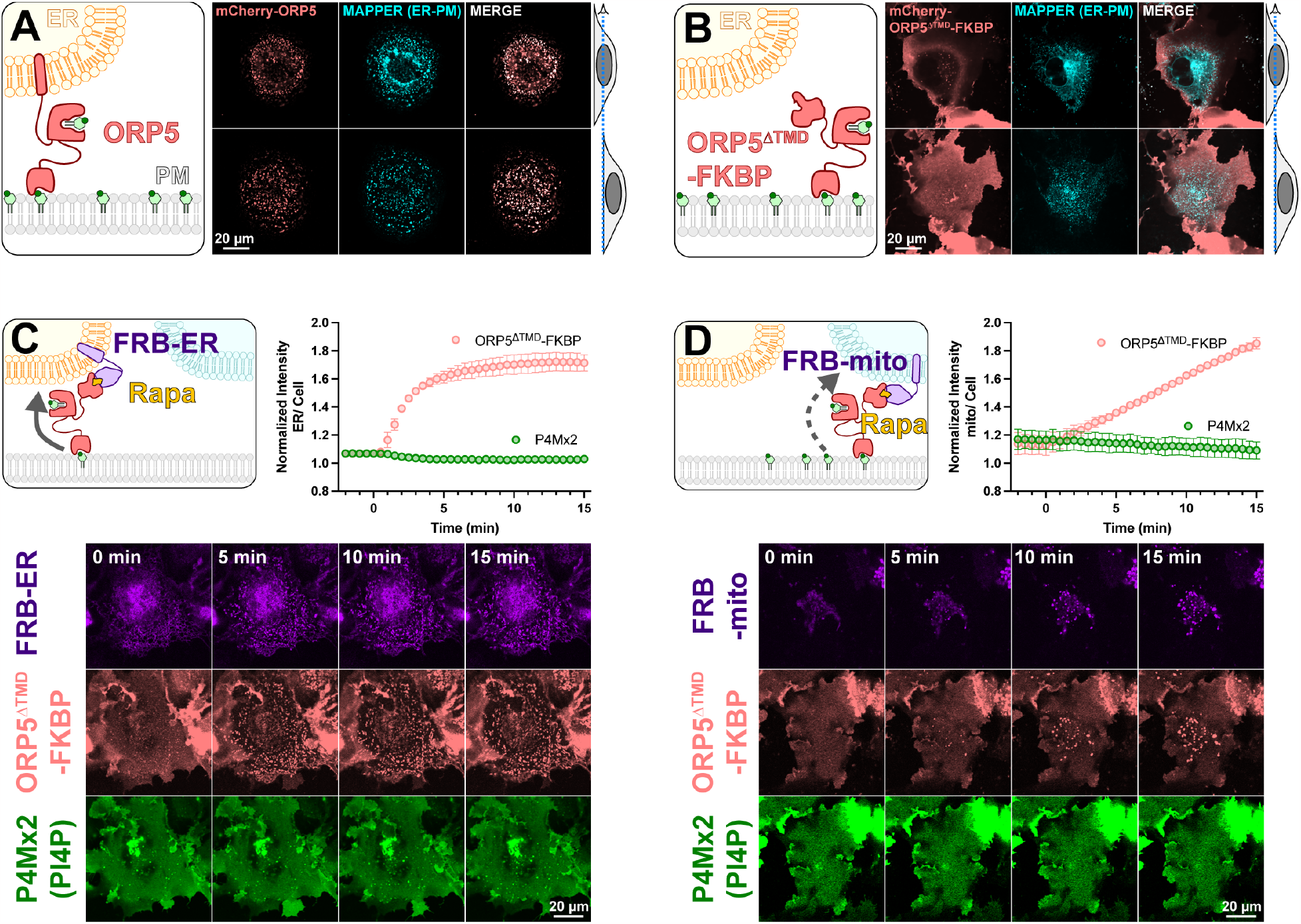
An engineered ORP5 ectopically targeted to PM-mitochondria contact sites fails to transfer PI4P. (**A**) mCherry-ORP5 localizes to ER-PM contact sites marked by the ER-PM contact site marker, MAPPER. Images show equatorial and ventral confocal sections of the same, representative cells. (**B**) Replacement of the ER-resident transmembrane domain (TMD) of ORP5 with FKBP leads to a delocalized PM localization, overlapping with but not restricted to MAPPER-labelled ER-PM contact sites. (**C**) mCherry-ORP5^ΔTMD^-FKBP can be returned to ER-PM contact sites by rapamycin-induced dimerization with an ER-localized FRB domain (iRFP-FRB-CYB5^tai^), though no enrichment of PI4P is seen at these induced contact sites. (**D**) mCherry-ORP5^ΔTMD^-FKBP can be targeted to ectopic mitochondrial-PM contact sites by rapamycin-induced dimerization with an mitochondria-localized FRB domain (AKAP1^N31^-FRB-iRFP), though no enrichment of PI4P is seen at these induced contact sites. For **C & D**, images show ventral confocal sections of representative cells; the graphs show the normalized mCherry-ORP5^ΔTMD^-FKBP or P4Mx2 PI4P biosensor enrichment at ER or mitochondrial contact sites, respectively. Data are grand means ± s.e. of three experiments, with 10 cells per experiment.

Previously, acute depletion of PM PI4P was demonstrated by replacing the N-terminal PH domain with the FK506 binding protein (FKBP) domain and inducing FKBP dimerization with a PM anchored FK506 and Rapamycin Binding domain of mTor (FRB) via the addition of rapamycin. This acute restoration of ORP5 ER-PM contact site localization was presumed to induce transport of PM PI4P to the ER, where it was degraded by SAC1 (Chung et al., 2015). To facilitate orthogonal contact site formation between the PM and another organelle (thereby uncoupling ORP5 transport from degradation by ER SAC1), we instead replaced the C-terminal ER-localizing helix with FKBP, a fusion henceforth termed ORP5^ΔTMD^-FKBP (cartoon in **Fig 1B**). This generates a PM localization no longer restricted to MAPPER-labelled ER-PM contact sites (**Fig. 1B**).

We then acutely restored ORP5^ΔTMD^-FKBP localization to ER-PM contact sites by rapamycin-induced dimerization with an ER-anchored FRB domain (**Fig 1C**). Such sites formed rapidly (completing within ∼5 min) when viewed by ventral confocal section. In parallel, we tracked the distribution of PI4P using the high affinity lipid biosensor EGFP-P4Mx2 (Hammond et al., 2014). As expected, we did not see accumulation of PI4P at these contact sites (see graph in **Fig. 1C**), presumably due to SAC1 activity in the ER. Rather, the fluorescence of PM PI4P seemed to decline, though this is not as obvious when viewed by confocal in such ventral focal planes.

We next sought to test whether orthogonal targeting of ORP5^ΔTMD^-FKBP to PM-mitochondrial contact sites would cause accumulation of PI4P in the latter organelle. Rapamycin induced dimerization recruited ORP5^ΔTMD^-FKBP to bright patches marked by mitochondria outer membrane-localized FRB, albeit somewhat more slowly than it recruits to ER-localized FRB (**Fig. 1D**). However, we saw no evidence of PI4P accumulation at these PM-mitochondria contacts (**Fig. 1D**).

Several scenarios could be envisioned that may preclude productive transfer of PM PI4P to mitochondria by ORP5^ΔTMD^-FKBP. One possibility is suggested by recent studies of the homologous yeast protein, Osh6p (Eisenreichova et al., 2021). That study reported that binding of PI4P by Osh6p is so tight that lipid transfer is inhibited. Hydrolysis of PI4P was required to deplete the lipid and permit PS binding in these experiments. We therefore hypothesized that SAC1 activity may in fact be required for productive release of the PI4P. To test this, we engineered SAC1 to ectopically target mitochondria. We replaced the ER-localized transmembrane domains with the mitochondrial outer membrane-targeting C-terminal 31 amino acids from Fis1 (Stojanovski et al., 2004), generating SAC1^mito^. This construct (fused to EGFP or TagBFP2) localized well to mitochondria (**Fig. 2A**), demonstrated by its close co-localization with a mitochondrial-matrix targeted mCherry (Filippin et al., 2005).

**Figure 2:**
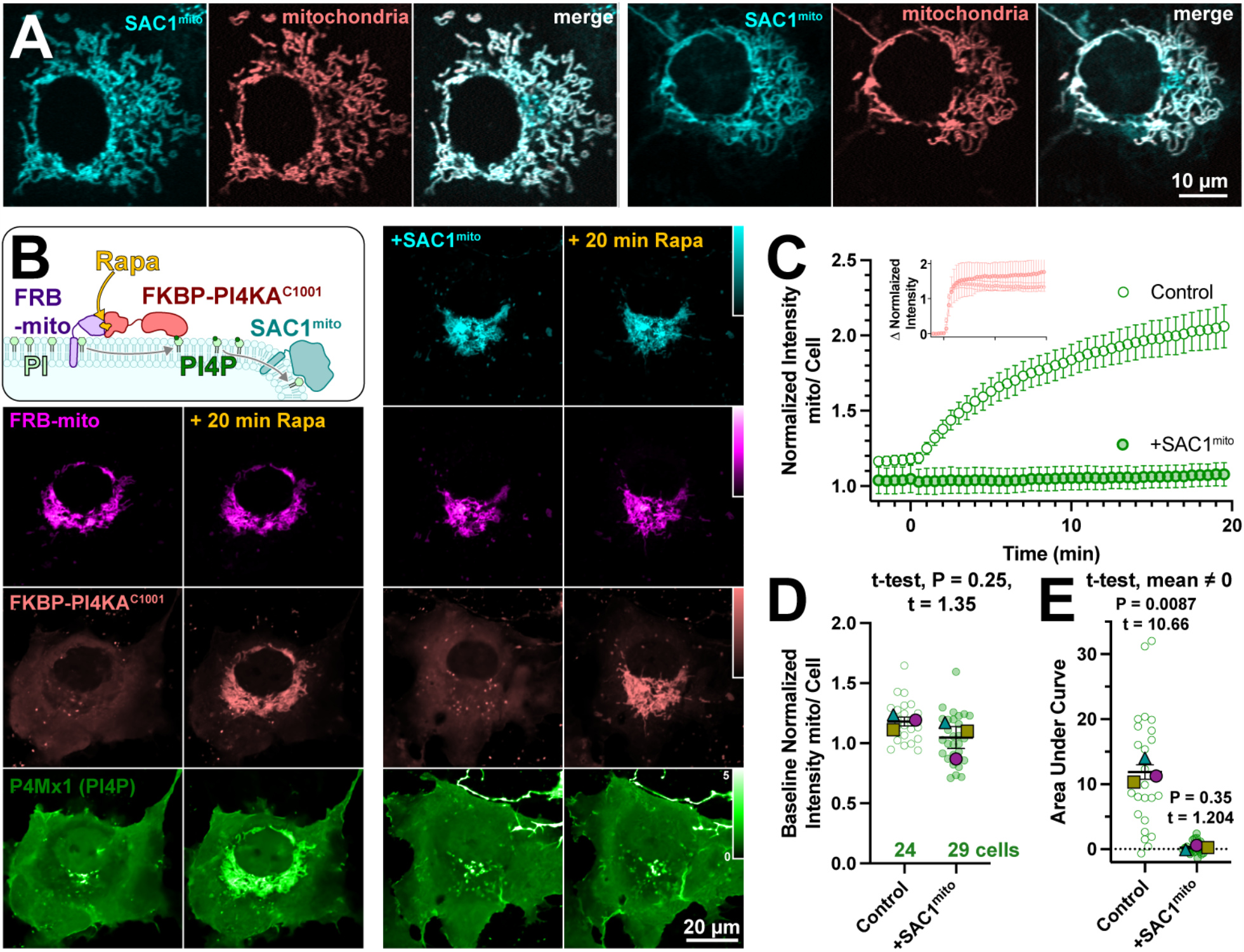
An engineered mitochondria-targeted SAC1 is active in cells. (**A**) SAC1^mito^ co-localizes with the mitochondrial marker, CAX8A^N29^x2-mCherry (pink). Images show representative equatorial confocal sections of two COS-7 cells. (**B**) Mitochondrial PI4P synthesis was induced by addition of rapamycin, which induces dimerization of over-expressed mCherry-FKBP-PI4K^C1001^ with iRFP-FRB-mito; this initiates ectopic PI4P synthesis on mitochondria outer membranes, detected with EGFP-P4Mx1 biosensor. The accumulation is blocked by co-expression of tagBFP2-SAC1^mito^. Images show equatorial confocal sections of COS-7 cells before or after addition of rapamycin. P4Mx1 intensity is normalized to the average cell intensity, as quantified in B. (**C**) PI4P accumulation on mitochondria is blocked by SAC1^mito^, despite equally efficient PI4KA^C1001^ recruitment (inset). (**D**) The slightly decreased baseline level of PI4P at mitochondria is not statistically significant by nested t-test. Smaller green points show individual cells (7-11 cells/experiment) and larger colored shaped show means of each of 4 experiments. Grand means ± s.e.m. are also indicated. (**E**) Complete reduction of mitochondrial PI4P synthesis, demonstrated by a grand mean of the area under the curve for SAC1^mito^-expressing cells of 0.24 ± 0.20 that is not significantly different from 0 by t-test.

To demonstrate activity of SAC1^mito^, we capitalized on our recent demonstration that PI4P synthesis can be ectopically induced on the mitochondria outer membrane by recruitment of the C-terminal helical-catalytic domain of PI4KA (Zewe et al., 2020). As shown in the left-hand panels of **Fig. 2B**, PI4P accumulation is readily detected with even a low affinity PI4P biosensor, P4Mx1 (Hammond et al., 2014). Co-expression of SAC1^mito^ under these conditions completely abolishes PI4P accumulation under these conditions (right hand panels in **Fig. 2B** and **Fig. 2C**), despite identical recruitment of the PI4K (inset in **Fig. 2C**). We did note a small decrease in baseline mitochondria associated P4M fluorescence in Sac1^mito^-expressing cells (**Fig. 1C**), though this was not statistically significant (**Fig. 2D**). The block in PI4P synthesis induced by SAC1^mito^ was complete, showing no significant deviation from 0 when measuring the area under the curve (**Fig. 2E**). These results show that we could reconstitute ER-like PI4P degradation on the mitochondrial outer membrane.

We next tested the ability of SAC1^mito^ to contribute to plasma membrane PI4P turnover. We accomplished this via inhibition of the major PM PI4P synthesizing enzyme, PI4KA, using the potent and selective inhibitor GSK-A1 (Bojjireddy et al., 2014). The rate of PI4P degradation or traffic from the PM can then be monitored by the drop in fluorescence of a high affinity PI4P biosensor, the P4C domain from *Legionella* SidC (Weber et al., 2014; Luo et al., 2015) in TIRFM. Note that we used P4C for many experiments as an alternative to the other high-affinity PI4P biosensor, P4Mx2. P4C has the advantage of not perturbing Golgi morphology at higher concentrations like P4Mx2 has been reported too (Hammond et al., 2014) – though both report the same changes in PM PI4P pools that we are concerned with here. We overexpressed both wild-type, ER-localized SAC1 together with SAC1^mito^, with either one being catalytically active whilst the other was an inactive (C389S) mutant (**Fig. 3A**). As previously reported (Sohn et al., 2016), over-expression of active, wild-type SAC1 accelerates PM PI4P depletion from the PM (**Fig. 3B**), presumably because the amount of SAC1 in the ER is rate-limiting for PM PI4P clearance. To our surprise, SAC1^mito^ also accelerated PM PI4P turnover (**Fig. 3B**) – at least as potently as the ER-localized, wild-type enzyme (**Fig. 3C**). To compare the relative activities of the enzymes, we transfected in either a low (50 ng) or high plasmid mass (500 ng) of either wild-type SAC1 or SAC1^mito^, and measured PI4P turnover in the same assay (**Fig. 3D**). We measured both the relative fluorescence intensity of the EGFP-tagged enzyme in TIRFM as a proxy for enzyme concentration proximal to the PM, and the rate of PM mCherry-P4Mx2 depletion. As shown in **Fig. 3E**, transfection of low or high plasmid mass led to the expected relative enzyme concentrations when viewed in TIRFM, with slightly increased concentrations of wild-type SAC1 compared to SAC1^mito^ at each plasmid mass – likely because there is more ER in proximity of the PM. Turning to PI4P turnover, both SAC1s caused a dose-dependent increase in PI4P turnover (**Fig. 3F**). SAC1^mito^ proved much more effective, however; transfection of a 50 ng concentration of SAC1^mito^ was much more potent than 50 ng of wild-type SAC1, and was even as effective as 500 ng wild type (**Fig. 3 F,G**).

**Figure 3:**
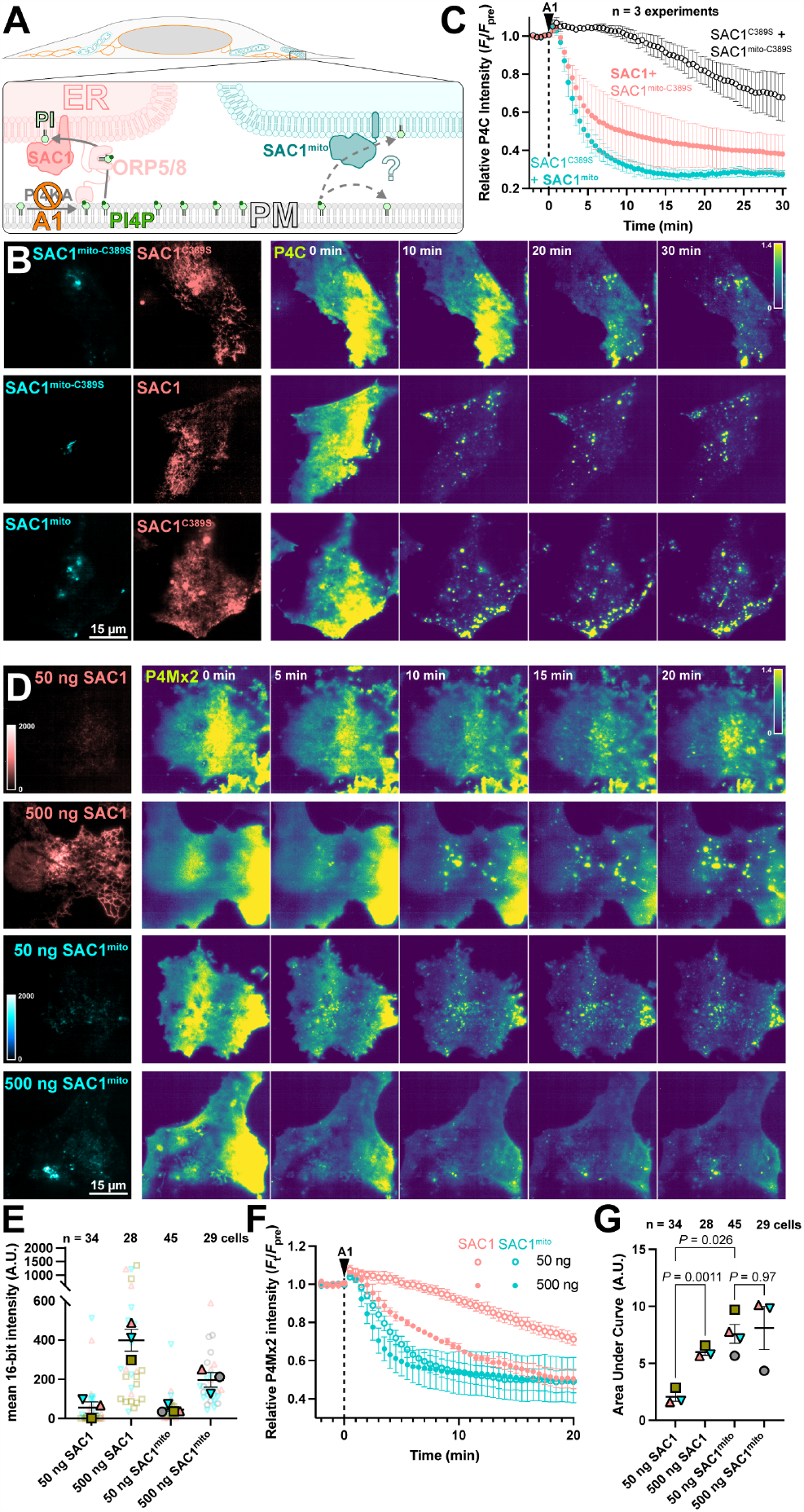
SAC1^mito^ accelerates PM PI4P turnover. (**A**) Native SAC1 in the ER is thought to turn over PM PI4P through transfer of the lipid from the PM to the ER; can SAC1^mito^ do the same? (**B-C**) SAC1^mito^ and over-expressed wild-type SAC1 accelerate PM PI4P turnover. (**B**) COS-7 cells were transfected with the PI4P biosensor EGFP-P4C-SidC, TagBFP2-SAC1^mito^ or its inactive C389S mutant, and mCherry-SAC1 or its C389S mutant. Cells were imaged in TIRF and 30 nM GSK-A1 added to inhibit PM PI4P synthesis at time 0. P4C fluorescence is normalized to the cell’s pre-bleach average as indicated. (**C**) Quantification of the P4C fluorescence data presented in B. Data are grand means ± s.e. of 3 independent experiments that measured 10-12 cells each. (**D-G**) SAC1^mito^ is more effective than over-expressed SAC1 at accelerating PM PI4P turnover. (**D**) COS-7 cells were transfected with the PI4P biosensor mCherry-P4M-SidMx2 and EGFP-SAC1^mito^ or EGFP-SAC1. Images of the EGFP were acquired with identical excitation and detection parameters and are displayed using the same intensity scale. Images are false colored for consistency with B-C. Cells were imaged in TIRF and 30 nM GSK-A1 added to inhibit PM PI4P synthesis at time 0. (**E**) Super-plot of the EGFP fluorescence intensity in the TIRF images; smaller empty symbols show individual cells; larger, filled symbols show the mean of individual experiments. Mean and s.e. of the 3-4 experiments is shown. (**F**) quantification of the P4Mx2 fluorescence intensity from D; data are grand means ± s.e. of 3-4 experiments. (**G**) Individual mean area under the curve analysis for individual experiments from F, with mean ± s.e. indicated. *P* values are from Tukey’s multiple comparisons test after a Mixed Effects model (*P* = 0.0089, F [DFn, DFd] = 30.75 [1.584, 3.168], Geisser-Greenhouse’s epsilon = 0.5279.

We previously reported that SAC1 is unable to directly hydrolyze PM PI4P from the ER in *trans* (Zewe et al., 2018). One potential explanation for enhanced PI4P turnover by SAC1^mito^ could therefore be *trans* activity of the engineered enzyme, conferred by the altered mitochondrially-targeted tail. Even at the greatly reduced area of PM-mito contact sites, SAC1^mito^ might clear PM PI4P much more efficiently than the ER-localized enzyme, which is reliant on lipid transfer. We explored this possibility with experiments to greatly expand the number of PM-mito contact sites in cells expressing SAC1^mito^; we aimed to test if PM PI4P turnover was further accelerated. This was accomplished through rapamycin-induced dimerization of PM localized FKBP and mitochondrially targeted FRB. As a control, we utilized a non-membrane targeted FKBP, which exhibited the accelerated PI4P turnover over 15 min (**Fig. 4A**). Recruitment of FRB-mito to FKBP tethered to the PM by extended flexible and helical linkers linked to the palmitoylated and myristoylated N-terminus of Lyn kinase greatly expanded PM-mito contact sites, but failed to further accelerate PM PI4P turnover (**Fig. 4B**). As an alternative approach, we utilized a deletion mutant of ORP5^ΔTMD^ missing the lipid transfer ORD (**Fig. 4C**); this produced more rapid formation of PM-mito contact sites, but also failed to accelerate PM PI4P clearance. An ORP5^TMD^-FKBP with a PI4P binding-deficient ORD behaved almost identically (**Fig. 4D**). Finally, we employed a PI4P-transfer competent ORP5^ΔTMD^-FKBP, but still saw no further acceleration of PM PI4P clearance (**Fig 4E**). Therefore, lipid transfer or simple membrane proximity did not appear to play a rate-limiting role in accelerating PM PI4P catabolism by SAC1^mito^.

**Figure 4:**
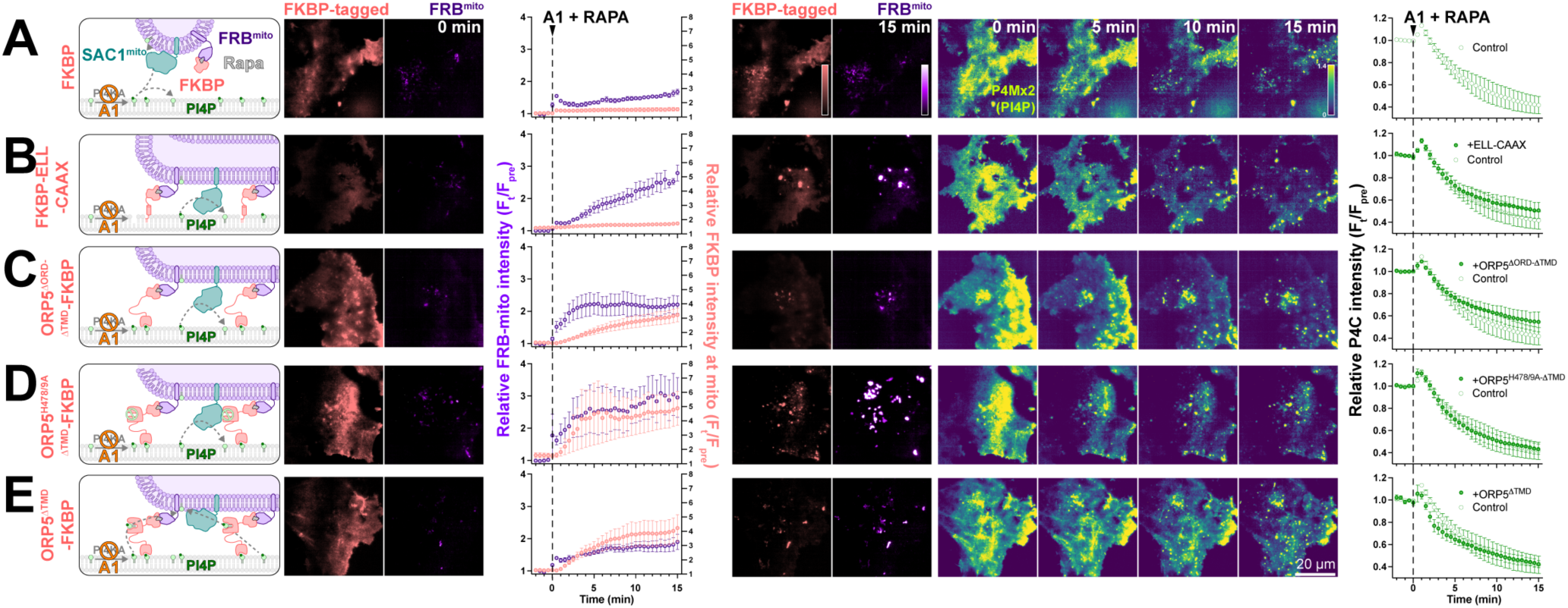
SAC1^mito^-mediated PM PI4P turnover is not enhanced at mito:PM contact sites. COS-7 cells were transfected with TagBFP2-Sac1^mito^, EGFP-P4C PI4P biosensor, iRFP713-FRB-Fis1^C31^ (FRB^mito^) and the indicated FKBP-tagged construct, After 18-24 h, they were subject to time lapse imaging by TIRFM. At time 0, 30 nM PI4KA inhibitor GSK-A1 was added together with 1 μM rapamycin to induced dimerization of FRB and FKBP. FKBP-tagged constructs were: (**A**) mCherry-FKBP as control; (**B**) an artificial PM-localized tethering construct, mCherry-FKBP-ELL-CAAX, consisting of the PM-anchored c-terminal tail of H-Ras linked to FKBP by an artificial helical and flexible linker mCherry-ORP5^ΔTMD^-FKBP; (**C**) a double deletion mutant of ORP5 lacking ER-localized transmembrane domain and the ORD, mCherry-ORP5^ΔORD-ΔTMD^-FKBP; (**D**) the PI4P-binding deficient mutant, mCherry-ORP5^H378/9A-ΔTMD^-FKBP; and (**E**) mCherry-ORP5^ΔTMD^-FKBP. For all panels, graphs show the ratio of mCherry-FKBP fluorescence at mitochondria to elsewhere in the TIRF footprint (i.e. PM), the relative intensity of the iRFP-FRB^mito^ signal and the total fluorescence intensity of EGFP-P4C in the TIRF footprint over time. Data are grand means ± s.e. of 3 experiments, analyzing 10-12 cells per experiment.

If transfer of PI4P at PM-mito contact sites or *trans* activity of SAC1^mito^ cannot explain the accelerated PM PI4P turnover, what could? Currently, PM PI4P turnover is thought to be driven by ORP5 and ORP8 at ER:PM contact sites (Chung et al., 2015; Sohn et al., 2018). However, we hypothesized that ORPs without such strict contact site localization may facilitate lipid transfer more broadly, perhaps presenting PI4P to our ectopically-targeted mitochondrial SAC1. Specifically, we studied ORPs 2, 3 and 6, since these paralogs have been reported to exhibit an at least partially cytosolic localization (Lehto et al., 2004; Wang et al., 2019a) and been implicated in PM phosphoinositide transfer cycles (Mochizuki et al., 2018; Wang et al., 2019a). When expressed in COS-7 cells, neonGreen-tagged ORP2, 3 and 6 all exhibited a primarily cytosolic localization, in contrast to the punctate, PM-apposed localization of ORP5 (**Fig. 5A**). Co-transfection of high-affinity PI4P (P4C) and PI(4,5)P_2_ (PH-PLCδ1) biosensors revealed much less prominent PM labelling by P4C in ORP2 and ORP5 transfected cells; in fact, PM PI4P was barely detectable in confocal (**Fig. 5A**). Quantitative image analysis confirmed reduced PM PI4P in ORP2 and ORP5-expressing cells (**Fig. 5B**), as previously shown for ORP5 (Sohn et al., 2018; Zewe et al., 2018; Chung et al., 2015). Although both ORP2 and ORP5 have been reported to transfer PI(4,5)P_2_ (Ghai et al., 2017; Wang et al., 2019a; Koponen et al., 2019), we could not detect a change in steady-state levels reported by PH-PLCδ1 (**Fig. 5C**). Co-expression of SAC1^mito^ had no additional effect on either PM PI4P or PI(4,5)P_2_ (**Figs. 5B**,**C**), although PM PI4P levels were so robustly depleted by ORP2 and ORP5 that additional effects of SAC1^mito^ would not be apparent in this assay.

**Figure 5:**
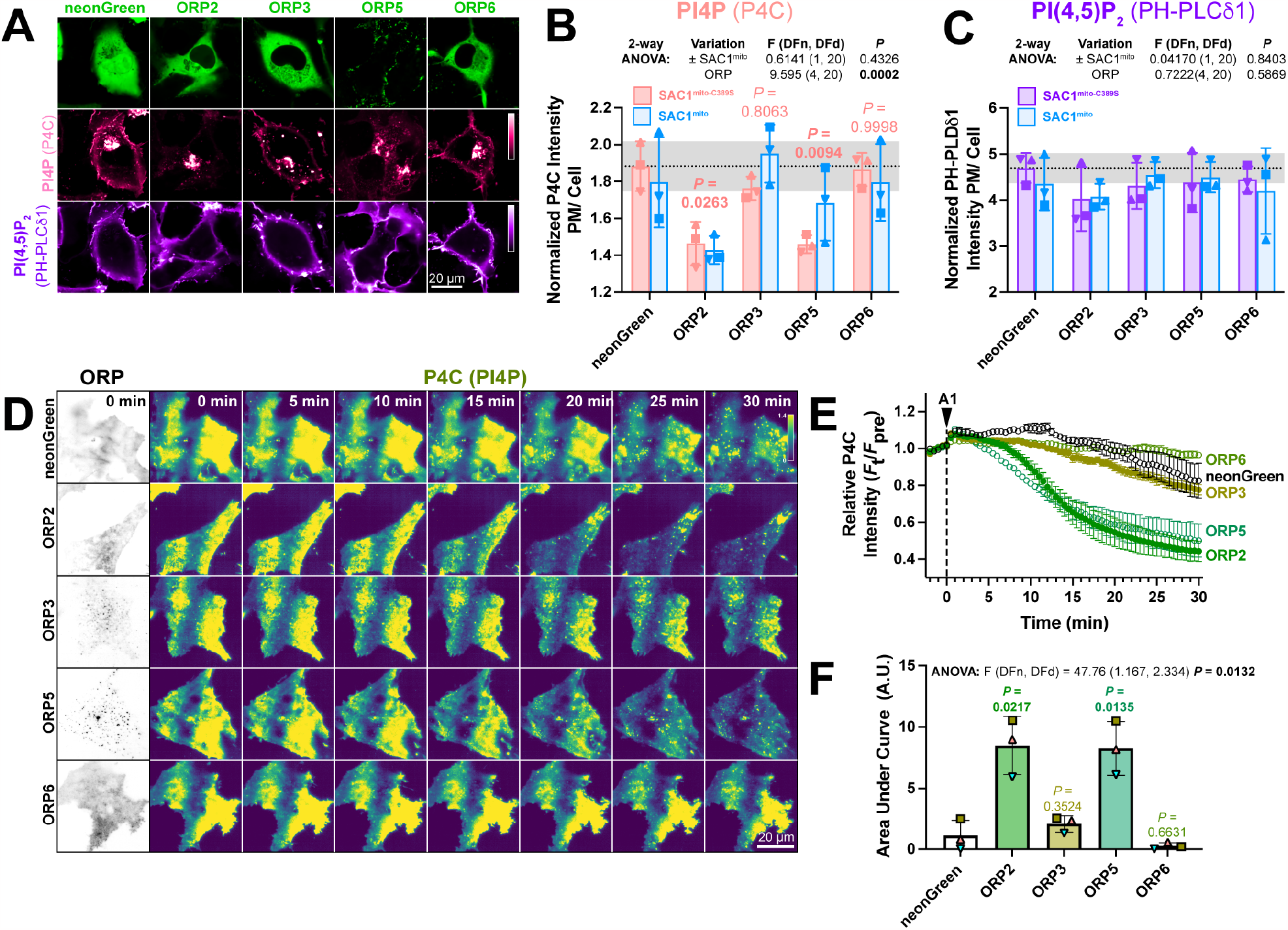
ORP2 over-expression accelerates PM PI4P turnover. (**A**) Representative confocal micrographs of COS-7 cells over-expressing the indicated neonGreen-tagged ORP construct (or neonGreen alone as control) together with the mCherry-P4C PI4P and iRFP713-PH-PLCδ1 PI(4,5)P_2_ biosensors and the inactive TagBFP2-SAC1^mito-C389S^ (not shown). (**B**) Quantification of the relative PM intensity of PI4P biosensor in COS-7 cells expressing SAC1mito-C389S (pink) or SAC1^mito^ (cyan); bars are grand means ± S.D. of three experiments, while symbols show the mean of 26-57 cells from each individual experiment. Results of a 2-way analysis of variance are shown above the graph; the *P* values in pink are from Sidak’s multiple comparison test, comparing the ^SAC1mito-^C389S-transfected experiments with the respective control group. (**C**) As B, but for the PI(4,5)P_2_ biosensor. (**D**) TIRFM images of COS-7 cells over-expressing the indicated neonGreen-ORP construct and the mRFP-P4C biosensor. Time lapse imaging of the P4C sensor is shown with the time since application of 30 nM GSK-A1 to inhibit PI4KA indicated. (**E**) Quantification of P4C fluorescence intensity shown in D; data are grand means of 3 experiments. (**F**) Quantification of the area under the curve from E. Bars are grand means ± S.D. of three experiments; data points are means of 5-12 cells from individual experiments. Results of a one-way ANOVA are indicated above the graphs, with *P* values above the graphs from a Dunnett’s multiple comparison test comparing with the control, neonGreen group.

We next tested effects of over-expressed ORPs on PM PI4P turnover by measuring PI4P clearance rates from the PM after inhibition of PI4KA. Consistent with previous reports (Sohn et al., 2018), we could see accelerated reduction of PI4P by ORP5 (**Figs. 5D, E**); of the other ORP proteins, only ORP2 showed accelerated clearance of remaining PM PI4P, with a similar rate to ORP5 (**Figs. 5D-F**). ORP3 and 6, by contrast, had no significant effect (**Figs. 5E**,**F**). Therefore, ORP2 appeared to be a viable candidate to facilitate transfer of PI4P from the PM, perhaps presenting it to ectopically expressed SAC1^mito^.

Despite its cytosolic localization when over-expressed (**Fig. 6A**), ORP2 contains a classical FFAT motif that binds ER-localized VAPA and VAPB proteins (Loewen et al., 2003). Therefore, the cytosolic localization of over-expressed ORP2 may be due to saturation of the endogenous VAP proteins, leaving no mechanism for ER targeting of excess neonGreen-ORP2. Indeed, co-expression of mCherry-VAPB caused ER localization of neonGreen-ORP2 (**Fig. 6B**). Nevertheless, ER localization of ORP2 was incomplete even with VAPB over-expression, showing significant cytosolic ORP2 “haze” between ER cisternae (**Fig. 6B**). We also expressed ORP2 at much lower levels for 4 hours, hoping to avoid saturation of endogenous VAP proteins. In this context, we still observed ORP2 being mainly cytosolic (**Fig. 6C**). Note, the faint puncta visible in the transfected cells are also visible in the neighboring untransfected cells, and represent autofluorescence that becomes visible at such low expression levels. Therefore, ORP2 does not appear to be tightly associated with the ER and can likely access any organellar cytosolic membrane leaflet.

**Figure 6:**
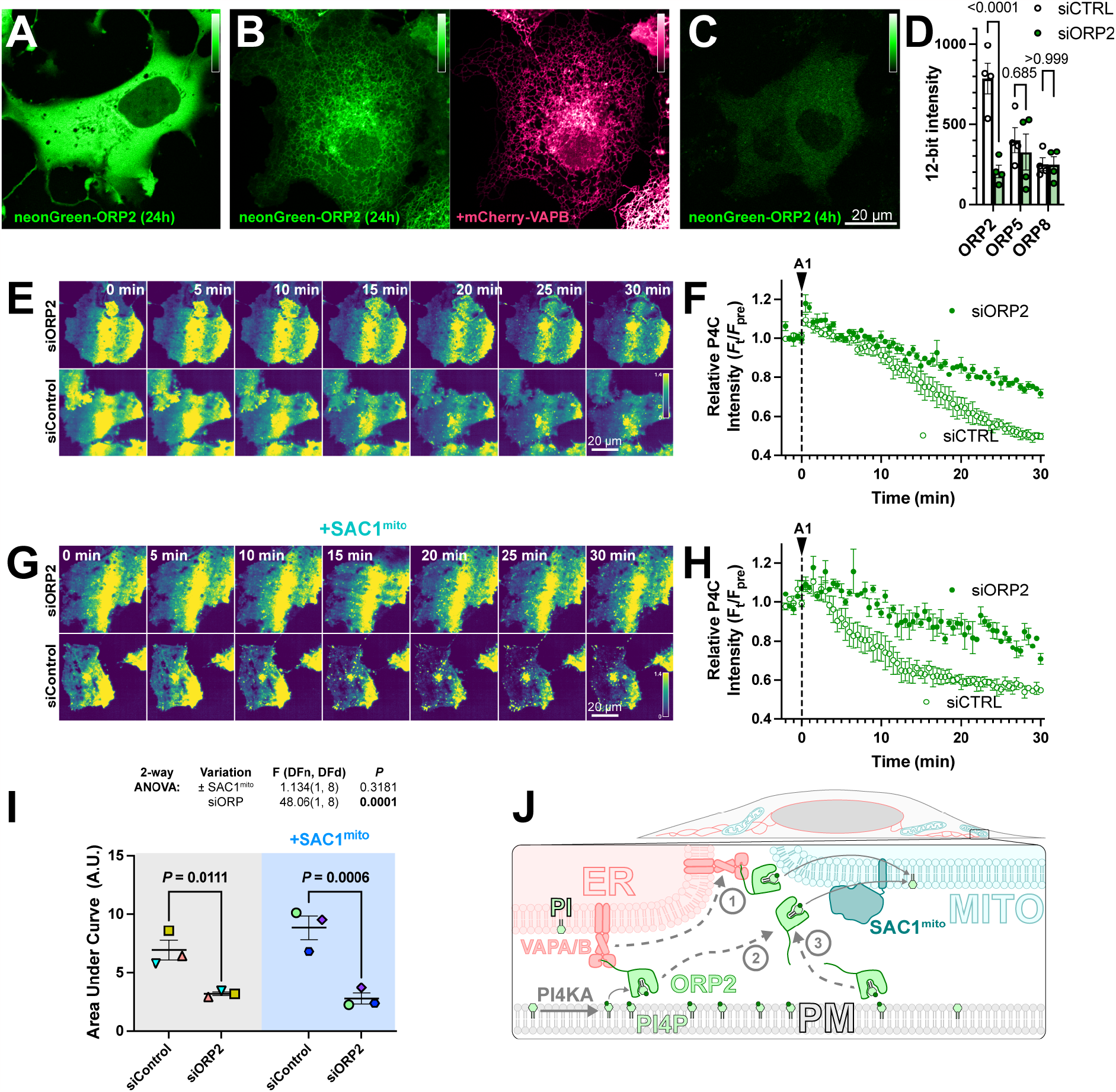
ORP2 is a major contributor to basal and SAC1^mito^-mediated PM PI4P turnover. (**A**) confocal section of a COS-7 cell over-expressing neonGreen-ORP2 for 24 h, showing a mainly cytosolic distribution. (**B**) as A, but the cell co-expresses mCherry VAPB, leading to partial localization of the neonGreen-ORP2 at the ER. (**C**) as A, but the cells were only transfected for 4 hours before imaging, leading to lower levels of fluorescence more consistent with endogenous expression levels. (**D**) TIRFM images of COS-7 cells expressing EGFP-P4C PI4P biosensor and either control or an ORP2-directed siRNA pool. Cells were treated with 30 nM GSK-A1 at time 0. (**E**) Quantification of the experiment depicted in D. Data are grand means ± s.e.m. of 3 experiments, quantifying 12 cells each. (**F-G**) as D-E, except cells were co-transfected with SAC1^Mito^. Data in G are grand means of 3 experiments, quantifying 10-12 cells each. (**H**) Area under the curve analysis of the data from E and G, with statistical analysis by two-way ANOVA. Pair-wise analysis gives the *P* values from a post-hoc Šídák’s multiple comparisons test. (**I**) Proposed model for ORP2 function in SAC1^mito^-mediated PM PI4P depletion; ORP2 removes PI4P from the PM, either tethered to VAPA or B at ER:PM contact sites (arrows **1** and **2**) or untethered anywhere in the PM (arrow **3**). From there, ORP2 diffuses to mitochondria to deliver PI4P for hydrolysis, either still tethered to VAPA or B (and thus occurring at ER:mitochondria contact sites, arrow **1**) or untethered from VAPs (arrow **3**).

We then sought to determine whether ORP2 contributes to PM PI4P turnover through small interfering RNA-mediated knock down of ORP2. siORP2 robustly depleted expression of neonGreen-ORP2, but had no effect on the other ORPs implicated in PM PI4P transfer, i.e. ORP5 or ORP8 **(Fig. 6D**). ORP2 depletion led to a significant slowing of PM PI4P loss (measured with P4C in TIRFM) after PI4K inhibition (**Fig. 6E, F**). Crucially, the knockdown also abolished the accelerated turnover of PI4P induced by SAC1^mito^ (**Fig. 6G, H** and **I**). Therefore, it seems highly likely that the accelerated PM PI4P turnover induced by SAC1^mito^ is because endogenous ORP2 transfers PI4P to the mitochondrial outer membrane for hydrolysis, a reaction that cannot happen in normally SAC1-devoid mitochondria (**Fig. 6J**).

## Discussion

To what extent does PM PI4P catabolism depend on lipid transport to the ER via membrane contact sites? We designed the experiments reported here to address this question. Whilst attempting to orthogonally target PM PI4P transport away from the ER to the mitochondrial outer membrane (**Figs. 1-2**), we made the unexpected discovery that PI4P turnover speeds up simply by targeting the major PI4P phosphatase, SAC1, to mitochondria (**Fig. 3**). No requirements for enhanced PM-mito contact sites were apparent (**Fig. 4**). This result was initially perplexing. Indeed, in a previous manuscript (now withdrawn), we erroneously attributed this enhanced turnover to an engineered ORP5 that we similarly targeted to mitochondria (Doyle et al., 2022). The experiments reported here show unequivocally that the orthogonally targeted ORP5 is not required (**Fig. 4**). Instead, we found that the non-contact site targeted OSBP related protein, ORP2, contributes to both basal PI4P turnover (**Figs. 5-6**) and the SAC1^mito^-induced accelerated turnover (**Fig. 6**). The data suggest that lipid transport, occurring separately from ER-PM membrane contact sites, could predominate the PI4P catabolic process. It also explains why native SAC1 is found throughout the ER, rather than enriched at ER:PM contact sites (Nemoto et al., 2000; Zewe et al., 2018).

It seems initially surprising that over-expression of SAC1 activity accelerates PI4P turnover at all: the fact that PI4P is never observed in the ER suggests that it is the rate of PI4P transport, rather than the rate of SAC1 hydrolysis, that is rate limiting. Yet, the acceleration of PI4P turnover reported here (**Fig. 3**) and by others (Sohn et al., 2018) after SAC1 over-expression suggests that SAC1’s catalytic activity is indeed rate limiting. So why isn’t PI4P accumulation observed in the ER? And if PI4P can be presented to and hydrolyzed by SAC1^mito^, why isn’t PI4P transfer to the mitochondrial outer membrane not normally observed? A clue comes from elegant *in vitro* work on Osh6 and ORP8 (Eisenreichova et al., 2021; Ikhlef et al., 2021); here, the ORD domain was shown to have exceptionally high affinity for PI4P, such that it inhibited binding and transport of the counter lipid, PS. These experiments predicted and demonstrated a requirement for PI4P hydrolysis for PS transport to occur. This high affinity suggests that the slow release (off rate) and rapid binding (on rate) of PI4P to ORD may cause rapid re-binding of released PI4P molecules at tethered membranes; lipid transfer would be effectively stalled, and the ORD would be saturated with PI4P. Efficient PI4P hydrolysis by SAC1 at the tethered membrane prevents this, facilitating counter lipid transport. Therefore, increasing SAC1 catalytic activity in the ER through over-expression effectively increases the off rate of the ORP and accelerates the whole transport cycle.

This mechanism also suggests why SAC1^mito^ is more effective than over-expressing wild-type SAC1: an ORP (like ORP2) not tethered to the PI4P replete membrane could bind any organelle membrane and deliver PI4P, either from ER contact sites (**Fig. 6I, steps 1 & 2**) or the cytosol (**step 3**). From here, the ORP will release but then rapidly rebind the PI4P until such time as it dissociates from the membrane. Only if the membrane has SAC1 activity is the PI4P not rebound (because it has been hydrolyzed). Therefore, we postulate that expression of SAC1^mito^ effectively harnesses the entire mitochondrial cytosolic membrane leaflet area as a novel sink for PI4P. This is far more effective than simply increasing the rate in the already available ER membrane surface area.

ORP2 has previously been reported to facilitate countertransport of PI(4,5)P_2_, as opposed to PI4P (Koponen et al., 2019; Takahashi et al., 2021; Wang et al., 2019a), which has also been reported for ORP5 and 8 in some studies (Ghai et al., 2017) but not others (Chung et al., 2015; Sohn et al., 2018; Ikhlef et al., 2021). We found much more profound effects of ORP2 on steady-state PI4P than PI(4,5)P_2_ (**Fig. 5**). Why the discrepancy? Yet another clue comes from the work on Osh6 and ORP8 (Eisenreichova et al., 2021; Ikhlef et al., 2021): here, ORD domains could bind both PI4P and PI(4,5)P_2_, but have a higher affinity for the former. Additionally, cells are known to preserve PI(4,5)P_2_ levels in the face of declining PI4P (Hammond et al., 2009, 2012; Bojjireddy et al., 2014). In fact, they even possess a homeostatic mechanism that will preserve PM PI(4,5)P_2_ levels at the expense of PI4P in the PM (Wills et al., 2022). Therefore, it is possible for an ORP to remove PI(4,5)P_2_, with or without PI4P, and still have a more profound impact on steady-state PI4P levels.

Overall, the most parsimonious explanation for our data is that ORPs such as ORP2 remove PI4P from the PM and present it to SAC1 for hydrolysis in the ER. The turnover number for ORPs is quite low, estimated to be approximately 0.5 transfer cycles per second (Lipp et al., 2019; Mesmin et al., 2013). Recent proteomic data estimated that there are approximately 36,000 copies of ORP2 in a HEK cell (Cho et al., 2022), giving an overall transfer rate of 18,000 PI4P per second. Given an estimated 10 million PI4P molecules per cell (Wills and Hammond, 2022), and assuming first order kinetics, the fractional rate constant, *k*, is 0.00036 s^−1^, giving a half-life (ln2/*k*) of 1925 s, or 32 min – not dissimilar to our observations in control cells (**Fig. 3C**). There is likely to be a minor contribution from ORP5 and 8 at the PM (Sohn et al., 2018), though this will decline as membrane targeting of these proteins depends on the continued presence of PI4P. Notably, neither knockdown of ORP5/8 or ORP2 has a complete block of PI4P turnover, underscoring contributions from all of these proteins.

Of course, ORP2 does not function in cells principally as a PI4P degrader. It, like other ORPs, is thought to harness the energy of the PI4P gradient for antiport of a second lipid (Arora et al., 2022). For ORP2, that lipid is cholesterol (Jansen et al., 2011). ORP2 has been shown to mediate the traffic of cholesterol between the ER and PM as well as endosomes (Takahashi et al., 2021; Koponen et al., 2019; Wang et al., 2019a). Clearly, there must be other determinants of membrane targeting for ORP2 to the PM as opposed to the Golgi, where OSBP predominates PI4P/cholesterol exchange (Mesmin et al., 2013, 2017). The fact that cholesterol is synthesized in the ER, and distributed to the PM from here, means that our ectopic targeting of SAC1 to mitochondria likely does not result in productive exchange cycles. Indeed, this is likely the key reason why physiologically speaking, the ER is the “sink” for PI4P transfer.

Given the advantages for speed and processivity of localizing lipid anti-port reactions to membrane contact sites, the strong effect of a non-contact site localized lipid transfer reaction that we report here seems counterintuitive. Are there any advantages to the cell of using such a non-constrained transfer mechanism? We can only speculate, but one possibility would be that it allows ORP2 to deliver PI4P, and hence retrieve cholesterol, from regions of the ER far removed from the PM. Given that cholesterol is often delivered to the ER from digested lipoproteins in the lysosomes rather than *de novo* synthesis, this may speed PM delivery: three-dimensional diffusion of ORP2 through the cytosol is likely to be much faster than the highly convoluted two-dimensional path cholesterol would have to take between lysosome:ER and ER:PM contact sites.

In conclusion, we discovered a surprising ability of SAC1 to enhance PM PI4P turnover, independent of the membrane it is targeted to – or the presence of membrane contact sites. The data implicate ORP2, and non-membrane contact site-targeted transfer proteins in general, in PM PI4P turnover.

## Materials and Methods

### Plasmids

The following plasmids were obtained as cited, or else generated by polymerase chain reaction and HiFi assembly (New England Biolabs) according to the manufacturer’s instructions:

**Table 1:**
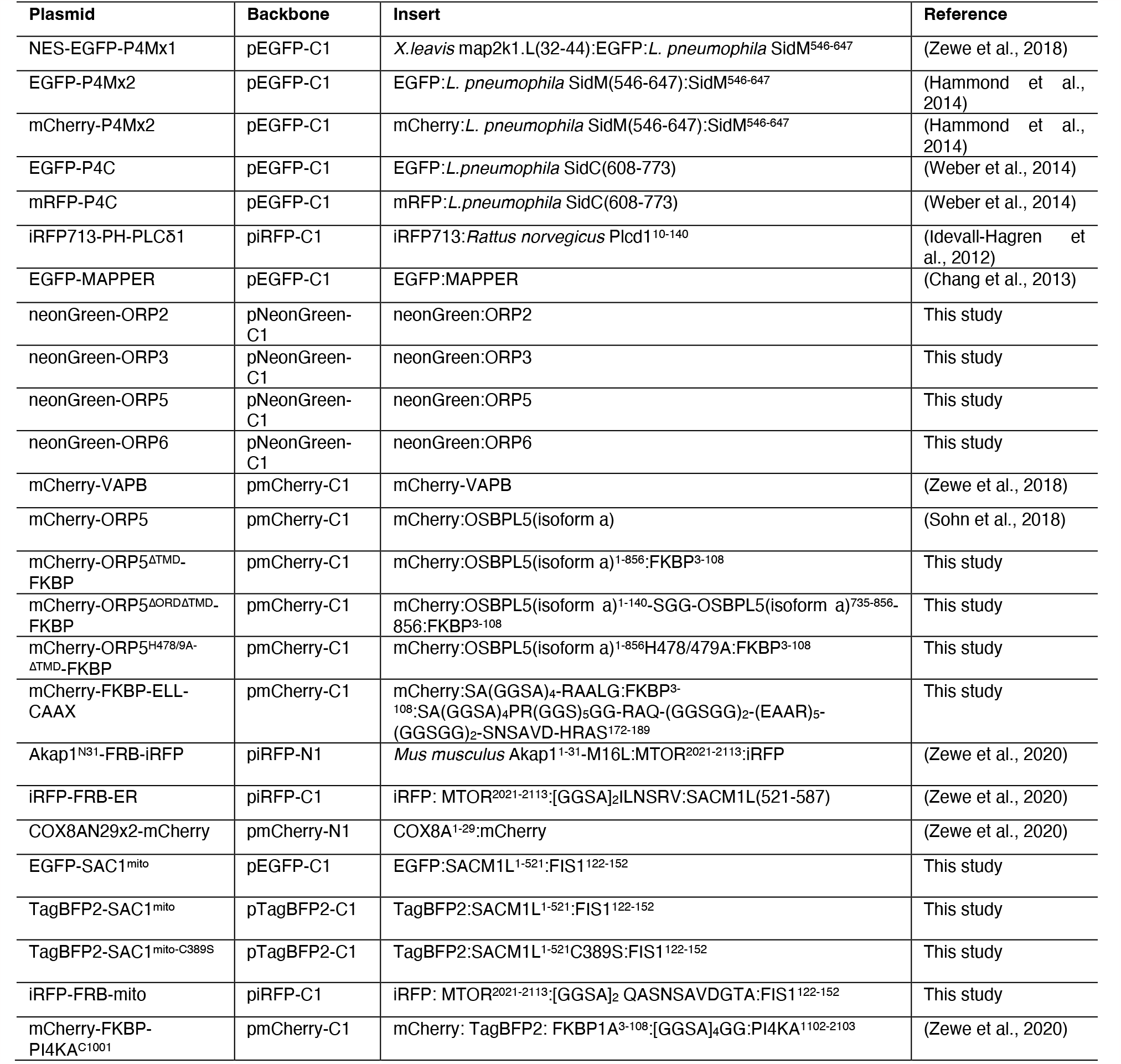
plasmids. HUGO Gene names and protein numbering are indicated. Human sequences were used unless a species is indicated (except for fluorescent proteins).

All plasmids were validated across the open reading frame by dideoxy sequencing.

### Cell Culture and Transfection

COS-7 cells (ATCC CRL-1651) were cultured in DMEM (5.56 mM glucose, glutaMAX supplement, Thermo Fisher 10567022) supplemented with 10% fetal bovine serum (ThermoFisher 10438-034), 10 u/ml penicillin and 10 μg/ml streptomycin (Thermofisher 15140122) and 1% (v/v) chemically defined lipid supplement (ThermoFisher 11905031). They were passaged twice weekly by rinsing in phosphate buffered saline and dissociation in Trypsin like-enzyme (ThermoFisher 12604039) before diluting 1:5 into fresh media. For transfection, cells were plated in 35 mm petri dishes with 20 mm aperture #1.5 optical glass bottoms (CellVis D35-20-1.5-N) coated with 5 μg/ml fibronectin (ThermoFisher 33016-015) at 25-50% confluence. Cells were transfected 1-24 h post seeding. For siRNA, 48 hours pre-cDNA transfection, cells were transfected with 50 pmol ORP2 smart pool (Horizon Discovery L-010361-01-0005) or siGENOME non-targeting pool (Horizon Discovery D-001206-13-05) precomplexed with 6.6 μl DharmaFECT (Horizon Discovery T-2001-01) in 400 μl serum-free medium before diluting to 2 ml for addition to cells (thus achieving a 25 nM final siRNA concetration). cDNA transfection was accomplished by the addition of 200 μl of Opti-MEM (ThermoFisher 51985091) pre-complexed for > 20 min with 1 μg DNA and 3 μg lipofectamine 2000 (ThermoFisher 11668019); DNA and lipofectamine were each diluted to 100 μl before combining. Specific plasmid mixtures are indicated in the figures. Prior to imaging, media was replaced with 2 ml of complete HEPES imaging media, composed of FluoroBrite media (ThermoFisher A1896702) supplemented with 10% fetal bovine serum, 1% (v/v) chemically defined lipid supplement, 2 mM glutaMAX (ThermoFisher 35050061) and 25 mM Na-HEPES, pH 7.4.

### Confocal Microscopy

A Nikon A1R-HD resonant scanning laser confocal microscope was used, mounted on a Nikon TiE inverted microscope stand. Images were collected through a 1.45 NA, plan apochromatic oil immersion objective (Nikon). Green (EGFP) and far red (iRFP713) fluorescence were co-excited using the 488 and 640 nm laser lines of a fiber-coupled LUN-V combiner, whereas blue (tagBFP2) and red (mCherry) fluorescence was excited on a subsequent line scan to avoid cross talk, using the 405 and 561 nm laser lines. Emission was collected using the following filters: blue (425-475 nm), green (505-550 nm), yellow-orange (570-620 nm) and far-red (650-850 nm). Confocal planes were collected with a pinhole of 1.2 Airy Units calculated for the far-red channel in resonant mode with 8 or 16 frame averaging. For time lapse imaging, up to 10 fields of cells were selected using the motorized stage and imaged every 30 s. Addition of rapamycin was accomplished by pipetting 0.5 ml of complete HEPES imaging media containing 5 μM rapamycin (Fisher Scientific BP2963-1) to the 2 ml already in the dish, achieving a final bath concentration of 1 μM.

### Total Internal Reflection Fluorescence Microscopy

A Nikon motorized TIRF illuminator mounted on a Nikon TiE inverted microscope stand was used. Laser excitation was fiber coupled from a four-line (405, 488, 561 and 638 nm) combiner (Oxxius). Emission was collected through dual pass filters from Chroma: blue/yellow-orange (420-480 nm / 570-620 nm) and green/far-red (505-550 nm / 650-850 nm). Imaging was performed through a 1.45 NA, plan apochromatic oil immersion objective (Nikon) using an Andor Zyla 5.5 sCMOS camera. For time-lapse imaging, up to 10 individual fields were marked using the motorized stage and imaged every 30 s. 0.5 ml of 5 μM Rapamycin (Fisher Scientific BP2963-1) + 150 nM GSK-A1 (Sigma SML2453) were added after 2 min to achieve bath concentrations of 1 μM and 30 nM, respectively.

### Quantitative Image Analysis and Statistics

For confocal image analysis, quantification of P4Mx2 signal at ER-PM or mitochondria-PM contact sites was accomplished by using the FRB-ER or mitochondria-FRB channels to generate a binary mask and measuring the normalized intensity of P4Mx2 inside this mask compared to outside. This and the auto thresholding procedure used to generate these masks has been outlined in detail in a recent protocols paper (Wills et al., 2021). Briefly, the FRB images were Gaussian filtered at lengths defined by 1 and 2 multiples of the far red-fluor airy disc size (for ER), or 1, 2, 3 and 4 times this size for mitochondria. Each filtered image was subtracted from the image filtered at next smallest length scale to generate wavelet images. These wavelets were multiplied together, and a threshold applied based on 0.5 standard deviations of the original image intensity, generating a binary mask. Finally, for mitochondria, the mask underwent a 2-pixel dilation to effectively cover all mitochondrial area in the image.

Analysis of TIRFM data was much simpler. After background subtraction, normalized intensity of the cell footprint was measured at each timepoint F_t_ and normalized to the mean of the pre-stimulation intensity (F_pre_).

Data analysis, statistics and graphs were plotted using Graphpad Prism 9. Details of statistical tests are given in the figure legends.

## Acknowledgements

We are grateful to Jen Liou (UT SouthWestern, USA) for sharing EGFP-MAPPER and Tamas Balla for sharing EGFP and mRFP-tagged P4C. We are grateful to all members of the Hammond lab for technical assistance with experiments during restrictive COVID mitigation. This works was supported by NIH grant R35GM119412. All authors declare no competing financial interests.

## References

Arora, A., J.H. Taskinen, and V.M. Olkkonen. 2022. Coordination of inter-organelle communication and lipid fluxes by OSBP-related proteins. Prog. Lipid Res. 86:101146. doi:10.1016/j.plipres.2022.101146.

Bojjireddy, N., J. Botyanszki, G. Hammond, D. Creech, R. Peterson, D.C. Kemp, M. Snead, R. Brown, A. Morrison, S. Wilson, S. Harrison, C. Moore, and T. Balla. 2014. Pharmacological and Genetic Targeting of the PI4KA Enzyme Reveals Its Important Role in Maintaining Plasma Membrane Phosphatidylinositol 4-Phosphate and Phosphatidylinositol 4,5-Bisphosphate Levels*. J Biol Chem. 289:6120–6132. doi:10.1074/jbc.m113.531426.

Chang, C.-L., T.-S. Hsieh, T.T. Yang, K.G. Rothberg, D.B. Azizoglu, E. Volk, J.-C. Liao, and J. Liou. 2013. Feedback Regulation of Receptor-Induced Ca2+ Signaling Mediated by E-Syt1 and Nir2 at Endoplasmic Reticulum-Plasma Membrane Junctions. Cell Reports. 5:813–825. doi:10.1016/j.celrep.2013.09.038.

Cheong, F.Y., V. Sharma, A. Blagoveshchenskaya, V.M.J. Oorschot, B. Brankatschk, J. Klumperman, H.H. Freeze, and P. Mayinger. 2010. Spatial Regulation of Golgi Phosphatidylinositol-4-Phosphate is Required for Enzyme Localization and Glycosylation Fidelity. Traffic. 11:1180–1190. doi:10.1111/j.1600-0854.2010.01092.x.

Cho, N.H., K.C. Cheveralls, A.-D. Brunner, K. Kim, A.C. Michaelis, P. Raghavan, H. Kobayashi, L. Savy, J.Y. Li, H. Canaj, J.Y.S. Kim, E.M. Stewart, C. Gnann, F. McCarthy, J.P. Cabrera, R.M. Brunetti, B.B. Chhun, G. Dingle, M.Y. Hein, B. Huang, S.B. Mehta, J.S. Weissman, R. Gómez-Sjöberg, D.N. Itzhak, L.A. Royer, M. Mann, and M.D. Leonetti. 2022. OpenCell: Endogenous tagging for the cartography of human cellular organization. Science. 375:eabi6983. doi:10.1126/science.abi6983.

Chung, J., F. Torta, K. Masai, L. Lucast, H. Czapla, L.B. Tanner, P. Narayanaswamy, M.R. Wenk, F. Nakatsu, and P.D. Camilli. 2015. PI4P/phosphatidylserine countertransport at ORP5- and ORP8-mediated ER–plasma membrane contacts. Science. 349:428–432. doi:10.1126/science.aab1370.

Doyle, C.P., L. Timple, and G.R.V. Hammond. 2022. Depletion of Plasma Membrane PI4P by ORP5 Requires Hydrolysis by SAC1 in Acceptor Membranes. Biorxiv. 2022.08.04.502856. doi:10.1101/2022.08.04.502856.

Eisenreichova, A., B. Różycki, E. Boura, and J. Humpolickova. 2021. Osh6 Revisited: Control of PS Transport by the Concerted Actions of PI4P and Sac1 Phosphatase. Frontiers Mol Biosci. 8:747601. doi:10.3389/fmolb.2021.747601.

Filippin, L., M.C. Abad, S. Gastaldello, P.J. Magalhães, D. Sandonà, and T. Pozzan. 2005. Improved strategies for the delivery of GFP-based Ca2+ sensors into the mitochondrial matrix. Cell Calcium. 37:129–136. doi:10.1016/j.ceca.2004.08.002.

Moser von Filseck, J. A. Čopič, V. Delfosse, S. Vanni, C.L. Jackson, W. Bourguet, and G. Drin. 2015. Phosphatidylserine transport by ORP/Osh proteins is driven by phosphatidylinositol 4-phosphate. Science. 349:432–436. doi:10.1126/science.aab1346.

Ghai, R., X. Du, H. Wang, J. Dong, C. Ferguson, A.J. Brown, R.G. Parton, J.-W. Wu, and H. Yang. 2017. ORP5 and ORP8 bind phosphatidylinositol-4, 5-biphosphate (PtdIns(4,5)P2) and regulate its level at the plasma membrane. Nat Commun. 8:757. doi:10.1038/s41467-017-00861-5.

Hammond, G.R.V., M.J. Fischer, K.E. Anderson, J. Holdich, A. Koteci, T. Balla, and R.F. Irvine. 2012. PI4P and PI(4,5)P2 Are Essential But Independent Lipid Determinants of Membrane Identity. Science. 337:727–730. doi:10.1126/science.1222483.

Hammond, G.R.V., M.P. Machner, and T. Balla. 2014. A novel probe for phosphatidylinositol 4-phosphate reveals multiple pools beyond the GolgiLocalization of PtdIns4P in living cells. J Cell Biology. 205:113–126. doi:10.1083/jcb.201312072.

Hammond, G.R.V., G. Schiavo, and R.F. Irvine. 2009. Immunocytochemical techniques reveal multiple, distinct cellular pools of PtdIns4P and PtdIns(4,5)P2. Biochem J. 422:23–35. doi:10.1042/bj20090428.

Idevall-Hagren, O., E.J. Dickson, B. Hille, D.K. Toomre, and P.D. Camilli. 2012. Optogenetic control of phosphoinositide metabolism. Proc National Acad Sci. 109:E2316–E2323. doi:10.1073/pnas.1211305109.

Ikhlef, S., N.-F. Lipp, V. Delfosse, N. Fuggetta, W. Bourguet, M. Magdeleine, and G. Drin. 2021. Functional analyses of phosphatidylserine/PI(4)P exchangers with diverse lipid species and membrane contexts reveal unanticipated rules on lipid transfer. BMC Biol. 19:248. doi:10.1186/s12915-021-01183-1.

Jansen, M., Y. Ohsaki, L.R. Rega, R. Bittman, V.M. Olkkonen, and E. Ikonen. 2011. Role of ORPs in Sterol Transport from Plasma Membrane to ER and Lipid Droplets in Mammalian Cells. Traffic. 12:218–231. doi:10.1111/j.1600-0854.2010.01142.x.

Koponen, A., A. Arora, K. Takahashi, H. Kentala, A.M. Kivelä, E. Jääskeläinen, J. Peränen, P. Somerharju, E. Ikonen, T. Viitala, and V.M. Olkkonen. 2019. ORP2 interacts with phosphoinositides and controls the subcellular distribution of cholesterol. Biochimie. 158:90–101. doi:10.1016/j.biochi.2018.12.013.

Lehto, M., J. Tienari, S. Lehtonen, E. Lehtonen, and V.M. Olkkonen. 2004. Subfamily III of mammalian oxysterol-binding protein (OSBP) homologues: the expression and intracellular localization of ORP3, ORP6, and ORP7. Cell Tissue Res. 315:39–57. doi:10.1007/s00441-003-0817-y.

Lipp, N.-F., R. Gautier, M. Magdeleine, M. Renard, V. Albanèse, A. Čopič, and G. Drin. 2019. An electrostatic switching mechanism to control the lipid transfer activity of Osh6p. Nat Commun. 10:3926. doi:10.1038/s41467-019-11780-y.

Loewen, C.J.R., A. Roy, and T.P. Levine. 2003. A conserved ER targeting motif in three families of lipid binding proteins and in Opi1p binds VAP. Embo J. 22:2025–2035. doi:10.1093/emboj/cdg201.

Luo, X., D.J. Wasilko, Y. Liu, J. Sun, X. Wu, Z.-Q. Luo, and Y. Mao. 2015. Structure of the Legionella Virulence Factor, SidC Reveals a Unique PI(4)P-Specific Binding Domain Essential for Its Targeting to the Bacterial Phagosome. PLoS Pathog. 11:e1004965. doi:10.1371/journal.ppat.1004965.

Mesmin, B., J. Bigay, J. Moser von Filseck, S. Lacas-Gervais, G. Drin, and B. Antonny. 2013. A Four-Step Cycle Driven by PI(4)P Hydrolysis Directs Sterol/PI(4)P Exchange by the ER-Golgi Tether OSBP. Cell. 155:830–843. doi:10.1016/j.cell.2013.09.056.

Mesmin, B., J. Bigay, J. Polidori, D. Jamecna, S. Lacas-Gervais, and B. Antonny. 2017. Sterol transfer, PI4P consumption, and control of membrane lipid order by endogenous OSBP. Embo J. 36:3156–3174. doi:10.15252/embj.201796687.

Mochizuki, S., H. Miki, R. Zhou, Y. Kido, W. Nishimura, M. Kikuchi, and Y. Noda. 2018. Oxysterol-binding protein-related protein (ORP) 6 localizes to the ER and ER-plasma membrane contact sites and is involved in the turnover of PI4P in cerebellar granule neurons. Exp Cell Res. 370:601–612. doi:10.1016/j.yexcr.2018.07.025.

Nemoto, Y., B.G. Kearns, M.R. Wenk, H. Chen, K. Mori, J.G. Alb, P.D. Camilli, and V.A. Bankaitis. 2000. Functional Characterization of a Mammalian Sac1 and Mutants Exhibiting Substrate-specific Defects in Phosphoinositide Phosphatase Activity*. J Biol Chem. 275:34293–34305. doi:10.1074/jbc.m003923200.

Prinz, W.A., A. Toulmay, and T. Balla. 2020. The functional universe of membrane contact sites. Nat Rev Mol Cell Bio. 21:7–24. doi:10.1038/s41580-019-0180-9.

Saint-Jean, M. de, V. Delfosse, D. Douguet, G. Chicanne, B. Payrastre, W. Bourguet, B. Antonny, and G. Drin. 2011. Osh4p exchanges sterols for phosphatidylinositol 4-phosphate between lipid bilayers. J Cell Biology. 195:965–978. doi:10.1083/jcb.201104062.

Sohn, M., P. Ivanova, H.A. Brown, D.J. Toth, P. Varnai, Y.J. Kim, and T. Balla. 2016. Lenz-Majewski mutations in PTDSS1 affect phosphatidylinositol 4-phosphate metabolism at ER-PM and ER-Golgi junctions. Proc National Acad Sci. 113:4314–4319. doi:10.1073/pnas.1525719113.

Sohn, M., M. Korzeniowski, J.P. Zewe, R.C. Wills, G.R.V. Hammond, J. Humpolickova, L. Vrzal, D. Chalupska, V. Veverka, G.D. Fairn, E. Boura, and T. Balla. 2018. PI(4,5)P2 controls plasma membrane PI4P and PS levels via ORP5/8 recruitment to ER–PM contact sitesRegulation of PM PI(4,5)P2 levels via ORP5/8. J Cell Biology. 217:1797–1813. doi:10.1083/jcb.201710095.

Stojanovski, D., O.S. Koutsopoulos, K. Okamoto, and M.T. Ryan. 2004. Levels of human Fis1 at the mitochondrial outer membrane regulate mitochondrial morphology. J Cell Sci. 117:1201–1210. doi:10.1242/jcs.01058.

Takahashi, K., K. Kanerva, L. Vanharanta, L. Almeida-Souza, D. Lietha, V.M. Olkkonen, and E. Ikonen. 2021. ORP2 couples LDL-cholesterol transport to FAK activation by endosomal cholesterol/PI(4,5)P2 exchange. Embo J. 40:e106871. doi:10.15252/embj.2020106871.

Wang, H., Q. Ma, Y. Qi, J. Dong, X. Du, J. Rae, J. Wang, W.-F. Wu, A.J. Brown, R.G. Parton, J.-W. Wu, and H. Yang. 2019a. ORP2 Delivers Cholesterol to the Plasma Membrane in Exchange for Phosphatidylinositol 4, 5-Bisphosphate (PI(4,5)P2). Mol Cell. 73:458–473.e7. doi:10.1016/j.molcel.2018.11.014.

Wang, Y., C.J. Mousley, M.G. Lete, and V.A. Bankaitis. 2019b. An equal opportunity collaboration between lipid metabolism and proteins in the control of membrane trafficking in the trans-Golgi and endosomal systems. Curr Opin Cell Biol. 59:58–72. doi:10.1016/j.ceb.2019.03.012.

Weber, S., M. Wagner, and H. Hilbi. 2014. Live-Cell Imaging of Phosphoinositide Dynamics and Membrane Architecture during Legionella Infection. Mbio. 5:e00839–13. doi:10.1128/mbio.00839-13.

Wills, R.C., C.P. Doyle, J.P. Zewe, J. Pacheco, S.D. Hansen, and G.R.V. Hammond. 2022. A Novel Homeostatic Mechanism Tunes PI(4,5)P2-dependent Signaling at the Plasma Membrane. Biorxiv. 2022.06.30.498262. doi:10.1101/2022.06.30.498262.

Wills, R.C., and G.R.V. Hammond. 2022. PI(4,5)P2: signaling the plasma membrane. Biochem J. 479:2311–2325. doi:10.1042/bcj20220445.

Wills, R.C., J. Pacheco, and G.R.V. Hammond. 2021. Phosphoinositides, Methods and Protocols. Methods Mol Biology. 2251:55–72. doi:10.1007/978-1-0716-1142-5_4.

Wong, L.H., A.T. Gatta, and T.P. Levine. 2019. Lipid transfer proteins: the lipid commute via shuttles, bridges and tubes. Nat Rev Mol Cell Bio. 20:85–101. doi:10.1038/s41580-018-0071-5.

Zewe, J.P., A.M. Miller, S. Sangappa, R.C. Wills, B.D. Goulden, and G.R.V. Hammond. 2020. Probing the subcellular distribution of phosphatidylinositol reveals a surprising lack at the plasma membrane. J Cell Biol. 219. doi:10.1083/jcb.201906127.

Zewe, J.P., R.C. Wills, S. Sangappa, B.D. Goulden, and G.R. Hammond. 2018. SAC1 degrades its lipid substrate PtdIns4P in the endoplasmic reticulum to maintain a steep chemical gradient with donor membranes. Elife. 7:e35588. doi:10.7554/elife.35588.

